# DNA Methylation by Restriction Modification Systems Affects the Global Transcriptome Profile in *Borrelia burgdorferi*

**DOI:** 10.1101/375352

**Authors:** Timothy Casselli, Yvonne Tourand, Adam Scheidegger, William K. Arnold, Anna Proulx, Brian Stevenson, Catherine A. Brissette

## Abstract

Prokaryote restriction modification (RM) systems serve to protect bacteria from potentially detrimental foreign DNA. Recent evidence suggests that DNA methylation by the methyltransferase (MTase) components of RM systems can also have effects on transcriptome profiles. The causative agent of Lyme disease, *Borrelia burgdorferi*, encodes two RM systems with N6-Methyladenosine (m6A) MTase activity. The specific recognition and/or methylation sequences have not been identified for either *B. burgdorferi* MTase, and it is not currently known whether these RM systems influence transcriptome profiles. In the current study, Single Molecule Real Time sequencing was utilized to map genome-wide m6A sites, and to identify consensus modified motifs in wild-type *B. burgdorferi* as well as isogenic MTase mutants. Four conserved m6A motifs were identified, and were fully attributable to the presence of specific MTases. Whole-genome transcriptome changes were observed in conjunction with the loss of MTase enzymes, indicating that DNA methylation by RM systems has effects on gene expression in *B. burgdorferi*. The results of this study provide a comprehensive view of the DNA methylation pattern in *B. burgdorferi*, and the accompanying gene expression profiles add to the emerging body of research on RM systems and gene regulation in bacteria.

**IMPORTANCE:** Lyme disease is the most prevalent vector-borne disease in North America, and is classified by the Centers for Disease Control and Prevention (CDC) as an emerging infectious disease with an expanding geographical area of occurrence. Previous studies have shown that the causative bacterium, *Borrelia burgdorferi*, methylates its genome using restriction modification systems that allow for the distinction of self from foreign DNA. Although much research has focused on the regulation of gene expression in *B. burgdorferi*, the effects of DNA methylation on gene regulation has not been evaluated. The current study characterizes the patterns of DNA methylation by restriction modification systems in *B. burgdorferi*, and evaluates the resulting effects on gene regulation in this important pathogen.

## INTRODUCTION

Lyme disease is the most prevalent vector-borne disease in North America (1, 2). Clinical manifestations can include arthritis, carditis, and neurological complications such as meningitis, cranial and peripheral neuritis, facial nerve palsy, and cognitive decline (3). This multisystem disease is caused by infection with a genetically heterogeneous group of spirochetes, including *Borrelia burgdorferi* and related species (4). Both inter- and intra-species phenotypic variation have been reported within this genospecies complex, and attempts have been made to correlate this diversity to genotypic differences (5-15). The contribution of alternative drivers of phenotypic diversity in *B. burgdorferi* such as epigenetic regulation of gene expression have not been evaluated.

DNA methylation is the product of methyltransferase enzymes (MTases), and has been described in both prokaryotes and eukaryotic organisms (16, 17). In prokaryotes, MTases are best understood as a component of restriction modification (RM) systems (18). RM systems are nearly ubiquitous in prokaryotes (19), and play a pivotal role in defense against foreign DNA. Specifically, type II RM systems encode both a MTase enzyme that recognizes and methylates a specific DNA sequence, as well as a restriction endonuclease (RE) enzyme that recognizes and cleaves at or near the unmethylated form of the recognition sequence (20). MTase and RE functions can be encoded on two separate proteins, or can serve as two distinct functional domains of a single protein (i.e. subtype IIC). As DNA methylation acts to protect from cleavage by the RE enzyme, the specific recognition motif is modified at ~100% of sites in the genome. In contrast to RM systems, orphan MTases that lack cognate RE enzymes are less common in prokaryotic genomes, and have roles in cell cycle regulation, phase variation, and regulation of gene expression (21, 22).

The predominant form of DNA methylation in prokaryotes is N6-Methyladenosine (m6A) (23), and two distinct m6A MTases have been described in *B. burgdorferi* (24, 25). Clones of the type strain B31 lacking plasmids lp25 or lp56 are much more competent when transformed with exogenous DNA but not *B. burgdorferi*-derived DNA, suggesting that these two genetic elements harbor RM systems (24, 25). Predicted amino acid sequence similarity implicated two genes encoding putative bifunctional MTase-RE proteins belonging to the PD…D/ExK superfamily of type IIC restriction enzymes; *bbe02* on lp25, and *bbq67* on lp56 (24). Inactivation of bbe02 replicates the transformation phenotype of lp25-minus strains, confirming this gene as the lp25-resident restriction endonuclease (26). Additionally, Southwestern blot analysis of *B. burgdorferi* genomic DNA shows decreased levels of m6A in strains lacking either bbe02 or lp56, supporting the bi-functional nature of these proteins (25).

Although the effects of bbe02 and bbq67 on transformation efficiency have been demonstrated, the specific recognition and/or methylation sequence motifs have not been identified for either *B. burgdorferi* MTase. It is not currently known whether the MTases associated with these RM systems can also function as gene expression regulators. The complement of identified MTases differs between commonly studied *B. burgdorferi* isolates (25), and mutant strains lacking endogenous MTases are often used as surrogates for “wild-type” strains in laboratory studies due to their increased transformation efficiency and ease of genetic manipulation (27-33). As such, a more comprehensive characterization of the DNA methylation systems in *B. burgdorferi* and their effects on gene regulation will aid in the interpretation of these studies, and could prove crucial for a more thorough understanding of *B. burgdorferi* biology.

In the current study, Single Molecule Real Time (SMRT) sequencing (34-36) was utilized to map the genome-wide DNA methylation pattern, and to identify consensus modified motifs in wild-type *B. burgdorferi* as well as isogenic MTase mutants. Four conserved m6A modification motifs were identified, and were fully attributable to the presence of either *bbe02* or lp56 (*bbq67*). Next, transcriptome profiles were compared between strains to determine the effects of altering global DNA methylation patterns on gene expression. Whole-genome transcriptome changes were observed in conjunction with the loss of MTase enzymes, indicating that DNA methylation by RM systems has effects on gene expression in *B. burgdorferi*. The results of this study provide a comprehensive view of the DNA methylation profile in *B. burgdorferi*, and the integrated gene expression profiles add to the relatively new body of research on gene expression consequences resulting from differentially methylated genomes by RM systems in bacteria.

## RESULTS AND DISCUSSION

### Single molecule real time sequencing reveals conserved non-palindromic m6A motifs

DNA methylation was assayed in genomic DNA isolated from *Borrelia burgdorferi* B31 (see Table S1 for strain information) using SMRT sequencing (Pacific Biosciences). Sequencing coverage varied from 151-fold to 428-fold among plasmids, with 969-fold coverage of the chromosome, and >99.9% concordance for all genomic elements compared to the published genome (Table S2; see Table S3 for reference genome accession numbers). In addition to base sequence information, the presence of DNA modifications including methylation are measurable using SMRT sequencing. The kinetic characteristics of DNA polymerization, such as the time duration between successive base incorporations (termed “interpulse duration”; IPD) are altered by the presence of a modified base in the DNA template. DNA methylation is characterized by detectable increases in the observed time lapse of base incorporation compared to expected (IPD ratio) (34-36). The phred-like modification quality value (QV) is assigned to each base using information on coverage and consistency of IPD ratios, and indicates the assay’s confidence in methylation calls. SMRT sequencing detected predominantly modified adenine bases in *B. burgdorferi*, as shown in a plot of modification QV against sequencing coverage (Figure 1A; red dots), matching the modification profile for m6A methylation. This finding is consistent with the predicted m6A MTase function of the RM systems in *B. burgdorferi*, as well as a previous report demonstrating the presence of m6A via Southwestern blot (25). Analysis of the sequences surrounding m6A sites revealed four conserved m6A modification motifs with a total of 5,606 m6A sites in the genome (Figure 1B). Only two of the motifs (CGRKA, GNAAYG) were modified at ~100% of available sites, consistent with RM protection from cleavage by a cognate RE enzyme. The remaining two motifs (DGDA<u>A</u>GG, DGGC<u>A</u>TG) were modified at 40% and 66% of available sites, respectively; a hallmark of orphan m6A modification not associated with restriction protection. This inefficient modification was reflected by a reduction in mean IPD ratios compared to fully methylated motifs (Figure 1D). Alignment of position weight matrices (Figure 1C) revealed conserved guanine residues at positions −3 and +2 surrounding the methylated adenine between the partially methylated DGDA<u>A</u>GG and DGGC<u>A</u>TG motifs and the fully methylated GNA<u>A</u>YG motif, suggesting these motifs may represent promiscuous methylation by a single MTase.

**Figure 1.**
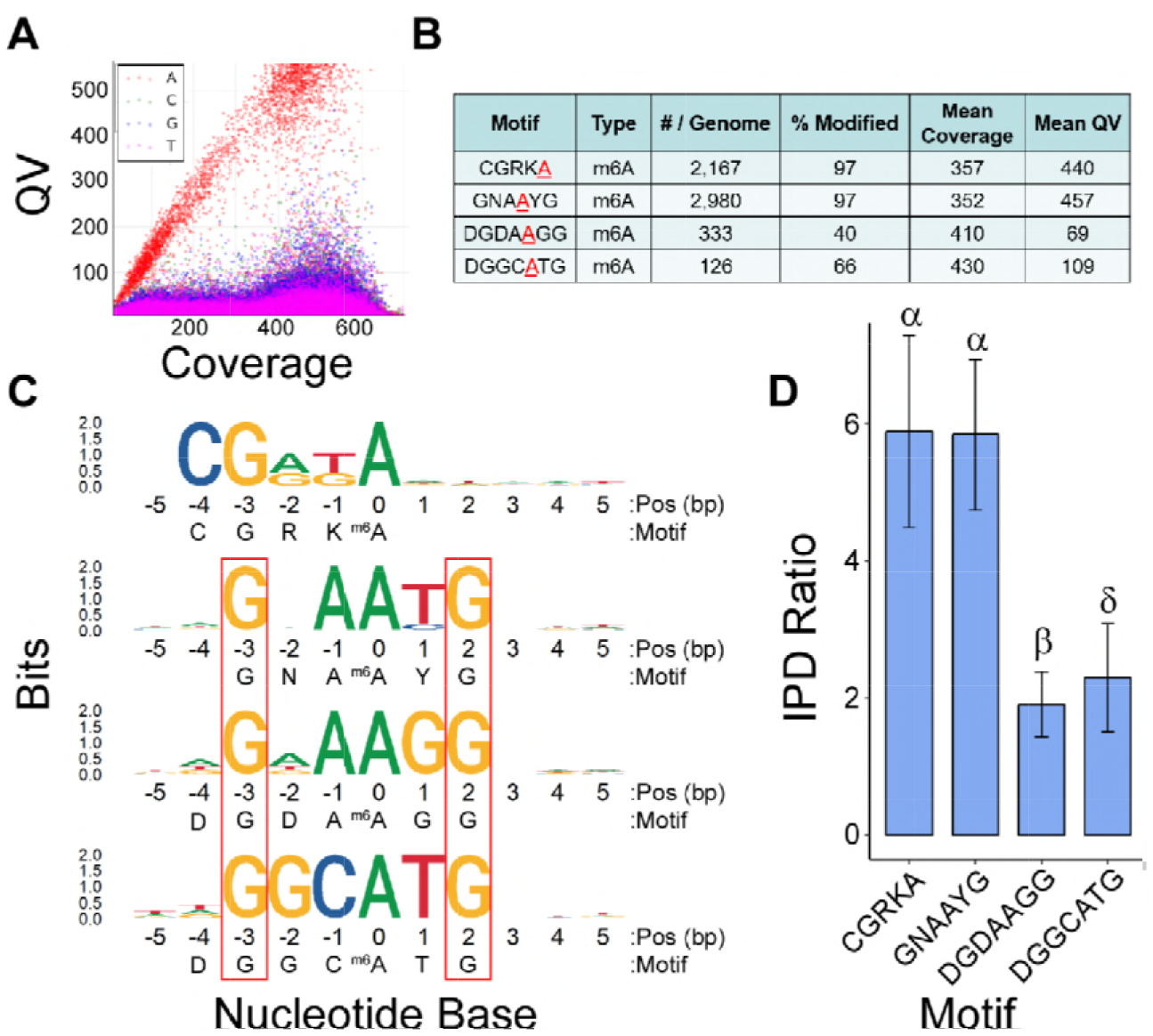
DNA modification in *B. burgdorferi* is predominantly m6A, and is located within conserved motifs. **A.** Dot plot demonstrating per-base modification score as a measure of sequencing coverage. Clustering of red dots shows adenine as the predominant modified base detected by SMRT sequencing in *B. burgdorferi*. **B.** Detected conserved m6A motifs along with the relative abundance and percent modified for each motif. Average sequencing coverage and modification scores are also shown. **C.** Sequence logo position weight matrices showing identified conserved modification motifs, with the modified adenine aligned at base position 0. Frequency of base occurrence at a given position is denoted by the height of each letter. Red boxes indicate conserved sites for three of the motifs, suggesting potential promiscuous modification by a single enzyme. **D.** Mean IPD ratios ± SD for all modified bases in each motif. Greek letters indicate significantly different groups as determined by one-way ANOVA followed by Holm-Sidak analysis of multiple comparisons (p<0.001). Decreased mean IPD ratios for DGDAAG and DGGCATG reflect incomplete modification at these sites.

All m6A motifs identified in *B. burgdorferi* were found to be non-palindromic (Figure 1), indicative of the recognition sequences of type IIS restriction endonucleases (e.g. FokI) (20). Type IIS REs cleave DNA at a specific location downstream of short, non-palindromic recognition sequences. Additionally, all motifs identified in this study were methylated on a single strand within the motif. While the majority of RM systems methylate both strands of the DNA molecule, a novel class of RM systems termed MmeI-family RM enzymes was recently found to possess type IIS restriction specificity, while the MTase function modifies a single strand within the recognition sequence (37, 38). This single-stranded methylation by MmeI-family enzymes is sufficient for restriction protection, likely due to the requirement for multiple unmethylated motifs for efficient cleavage.

### Methylated adenines are not evenly distributed throughout the *B. burgdorferi* genome

In order to evaluate the distribution of m6A sites throughout the genome, the number of modification motifs were determined per 1000 bp region. As shown in Figure 2A-B, there was variability in the frequency of m6A sites both between and within genomic elements. The chromosome had an increased median number of m6A per 1000 bp compared to plasmids with the exception of cp26 and lp28-2, and the median number per region also differed between different plasmids as determined by non-overlapping 95% confidence intervals. Each genomic element typically contained at least one outlier region of higher or lower frequency of modifications. Interestingly, the center of the chromosome contained a cluster of increased m6A outliers (Figure 2B) that mapped closely to the reversal in GC skew (data not shown), suggesting a possible role in DNA replication (39); however this pattern was not consistent for any of the plasmids presumed to have similar replication mechanisms. Strikingly, the plasmid lp21 contained just four conserved m6A motifs (Figure 2A), all within the first 3,074 bp of the “left” end (Figure 2B). This plasmid contains a 63 bp tandem repeat element that spans ~11kb (40). As the repeated element does not contain any conserved sites for m6A modification, lp21 contains very few modified bases overall.

**Figure 2.**
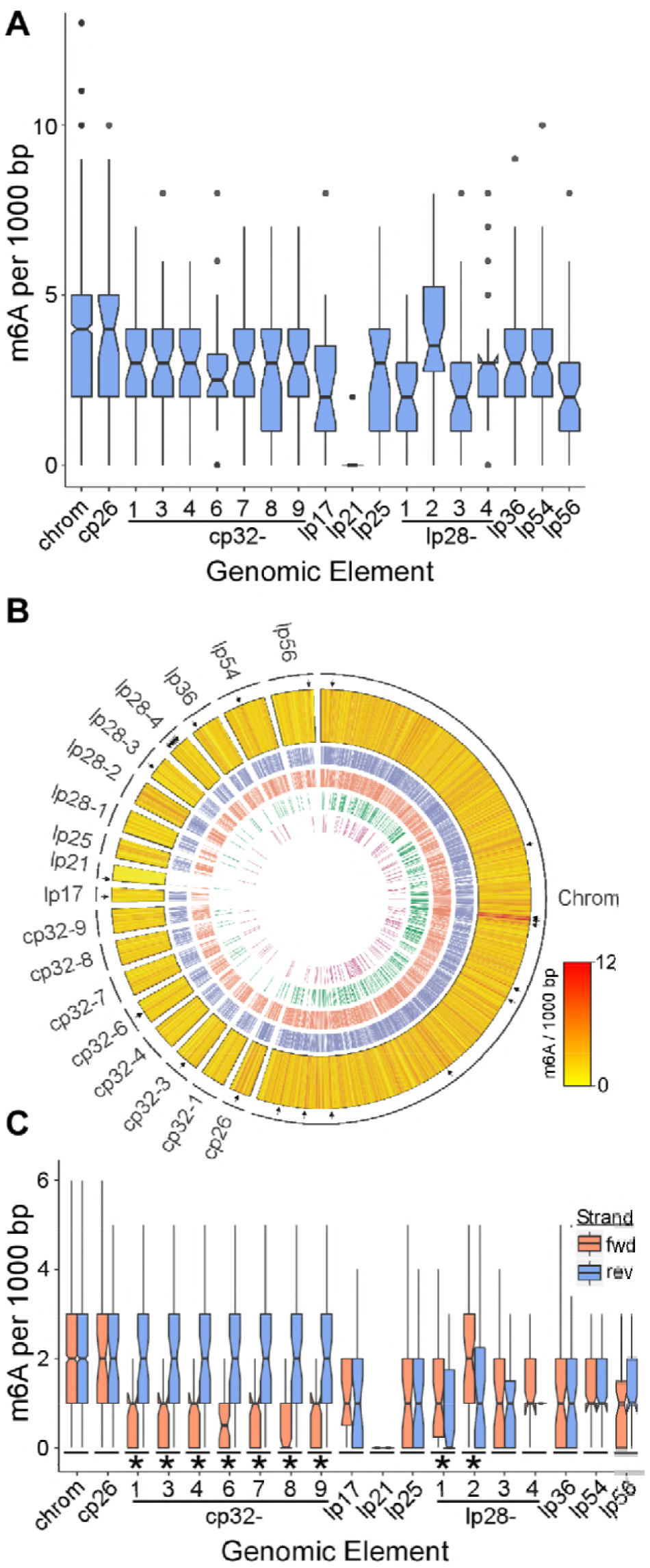
m6A distribution is not uniform across the genome. **A.** Distribution of total m6A sites per 1000 bp for each genomic element. Notches represent 95% CI of the median as determined by median ± 1.58*IQR/(n^½^). Non-overlapping CIs demonstrate variation in methylation frequency between elements. Dots represent outlier regions within each element. **B.** Circos plot showing heatmap of modifications per 1000 bp (yellow-red; outer track). Scale bar indicates number of m6A per region. Black tick marks outside heat map show locations of outlier regions from (A). Inner tracks show individual motif locations (GNAAYG, blue; CGRKA, red; DGDAAGG, green, DGGCATG, purple). **C.** Per-strand distribution of m6A sites per 1000 bp for each genomic element. Notches represent 95% CI of the median. Asterisks below boxes represent genomic elements with non-overlapping 95% CIs between forward and reverse strands. Note that outliers are not displayed.

Methylation of all four identified m6A motifs was found to be localized to a single strand within the motif. Plotting m6A per 1000 bp fragment by strand (forward vs reverse) revealed a strong reverse-strand bias for all of the cp32 plasmids, as well as less striking forward-strand biases for lp28-1 and lp28-2. The cp32 plasmids are stable prophages (41-43) that contain an average strand bias for encoded genes of ~15:1 not seen in the rest of the genome (i.e. chromosome coding strand bias of 1:1), suggesting a possible coding-strand bias for m6A motifs. Separate analyses of the distributions of forward/reverse strand m6A sites did not reveal any obvious patterns with implications for DNA replication (i.e. leading/lagging strand bias; data not shown).

Of the 1,784 genes and ncRNAs annotated or reported in *B. burgdorferi* (44) (Table S3), 76% contained at least one m6A motif within the gene body (Table 1; see Table S4 for complete gene body m6A counts). The median number of m6A sites per gene was three, with a maximum of 33. Examination of 200 bp regions upstream of these genes that may contain promoter and/or 5’-UTR elements revealed 46% of these upstream regions possessing at least one modification motif, with a median of one and a maximum of 7 (Table 1; see Table S5 for complete upstream region m6A counts).

**Table 1.** Number of *B. burgdorferi* genes containing m6A motifs

|  | Modified /<br>Total (%)<sup>a</sup> | Median m6A /<br>Gene (95% CI) | Maximum<br>m6A / gene |
| --- | --- | --- | --- |
| Gene Bodies | 1349 / 1784 (76%) | 3 (2.83-3.17) | 33 |
| Upstream region<sup>b</sup> | 758 / 1784 (42%) | 1 (0.94-1.06) | 7 |

<sup>a</sup>Number of regions containing at least one m6A motif.
<sup>b</sup>Modifications found in 200 bp region located 5' of start site

All four conserved motifs showed a significant bias towards the sense coding strand of genes when the total number of gene-body m6A sites were considered (p<0.002; Figure 3A), whereas the GNA<u>A</u>YG motif appeared to contribute most to this bias on a per-gene basis (Figure 3B). This bias toward the sense strand may be a mechanism to maximize genomic DNA methylation events, while minimizing the effects on gene expression. Modified bases on the template DNA strand (i.e. the anti-sense strand) may slow the rate of transcription elongation, much like the premise for SMRT sequencing described for DNA polymerase. This has been described for various DNA modifications including m6A in eukaryotic systems (45, 46), however the effects of m6A modification on prokaryote RNA polymerase kinetics is not known. Nonetheless, it is possible that overrepresentation of m6A modifications on the sense strand of genes may minimize any potential effects on RNA polymerase kinetics in *B. burgdorferi*.

**Figure 3.**
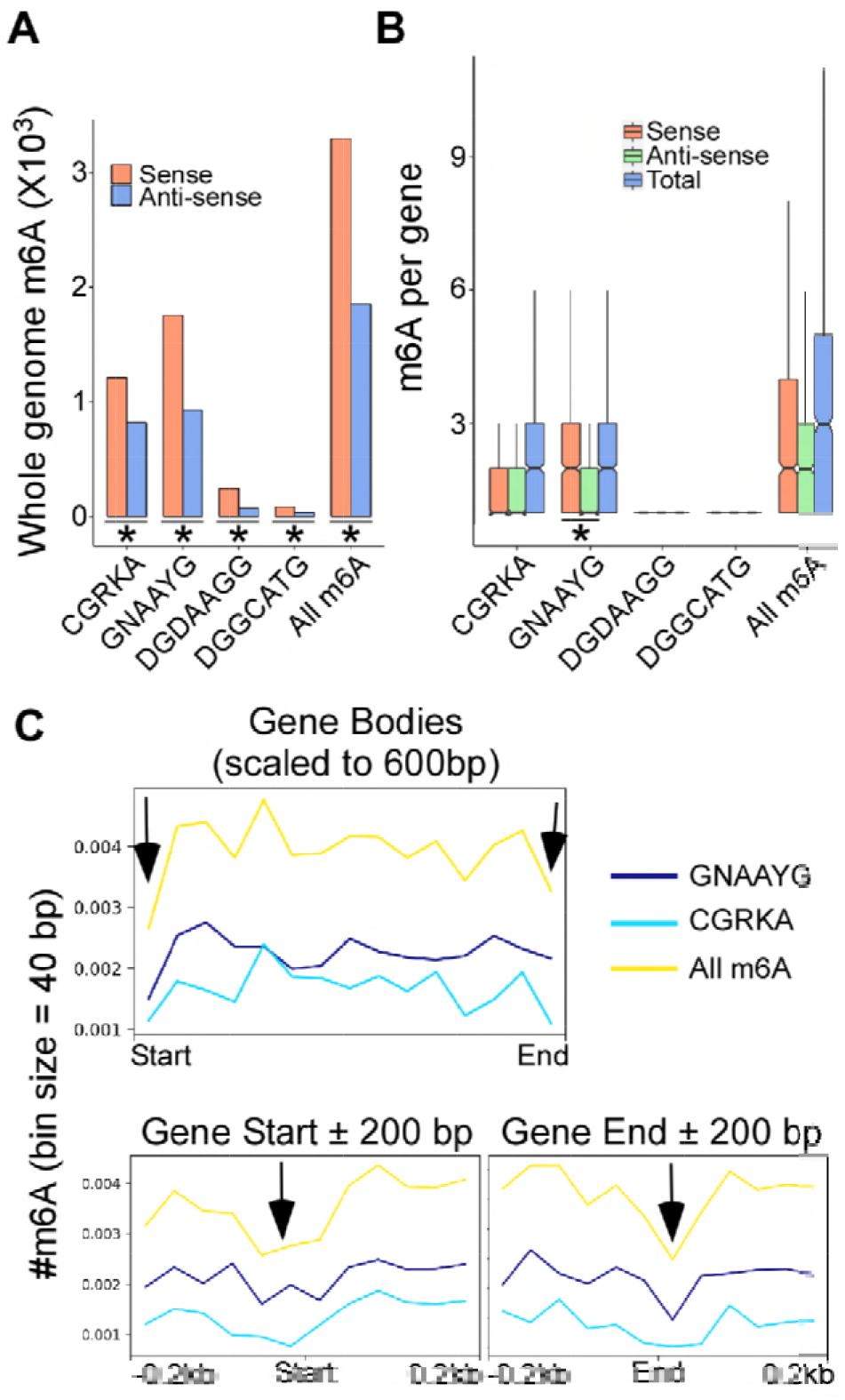
Gene body m6A shows a coding strand bias. **A.** Total gene body m6A counts by coding strand (sense vs. anti-sense). All motifs showed a significant sense strand bias as determined by Fisher’s Exact test (p<0.002), as denoted by asterisks below bars. **B.** Distribution of m6A counts per gene by coding strand. Asterisk indicates a sense strand bias for the GNAAYG motif as determined by non-overlapping CIs between sense and anti-sense distributions. Outliers are not displayed. **C.** Metagene analysis showing average m6A per 40 bp bin size across all gene bodies scaled to 600 bp (top panel), as well as at gene start and end sites ± 200bp (bottom panels). Black arrows show an apparent reduction in m6A frequency near the start and end coordinates.

Hotspots of DNA modifications surrounding and within gene bodies can provide clues as to potential functional consequences. Metagene analysis of m6A distributions within genes did not reveal any obvious bias of m6A sites towards the 5’ or 3’ end of the genes or clustering at putative promoter elements, however there was an apparent decrease in m6A frequency at both the gene start and end sites (Figure 3C). Unsupervised clustering using different parameters did not reveal any reliably reproducible sub-groups with patterns of m6A enrichment across gene bodies or at the gene start/end sites (data not shown).

### m6A motifs detected by SMRT sequencing are attributable to *bbe02* and *bbq67*

The genome sequence of *B. burgdorferi* encodes three intact putative bifunctional MTase-RE proteins belonging to the PD…D/ExK superfamily of type IIC restriction enzymes; *bbe02* on lp25, *bbq67* on lp56, and *bbh09* on lp28-3 (24, 47). Of these, only *bbe02* and *bbq67* have been demonstrated to contribute to restriction protection against exogenous DNA, and to the presence of genomic m6A (24, 25). In order to assign recognition sequence specificity to individual MTases, the methylomes of mutant strains lacking *bbe02* alone or both *bbe02* and *bbq67* were assayed using SMRT sequencing (see Table S1 for strain information).

The *B. burgdorferi* B31 clonal isolate 5A18 was previously characterized to be lacking the linear plasmid lp56 harboring the *bbq67* gene (48). An isogenic mutant of 5A18 (5A18-NP1; further referred to as BbΔe02Δq67) was also previously generated to contain an insertional inactivation in the *bbe02* gene using a kanamycin resistance cassette (*kan*) (26). In order to generate a mutant strain lacking *bbe02* alone, the NP1 mutation was introduced into wild-type *B. burgdorferi* as described in Materials and Methods (A3-NP1; further referred to as BbΔe02). Wild-type and mutant strains were screened by PCR for the presence of MTase genes (Figure 4A; see Table S8 for primer sequences). As expected, template DNA from BbΔe02Δq67 did not produce a PCR product using primers specific for lp56 harboring the *bbq67* gene. In contrast, all strains were positive by PCR screening using primers specific for the deleted region of *bbe02*. As *bbh09* and *bbe02* are homologues with 89% nucleotide identity (47), this may have been the result of non-specific amplification of *bbh09*. To resolve this issue, NheI-digested plasmid DNA was analyzed by Southern blot and probed for the deleted region of *bbe02* as well as the *kan* insertion marker (Figure 4B). As expected, only DNA from the wild-type strain was positive for *bbe02* and negative for the *kan* marker, whereas both BbΔe02 and BbΔe02Δq67 were negative for *bbe02* and positive for *kan*. Additionally, a positive band was seen for all three strains at the expected size for the fragment containing the *bbh09* gene when probed for *bbe02*. Collectively, these data confirm the specific deletion of *bbe02* in both mutants, without disruption of the the homologous *bbh09* locus.

**Figure 4.**
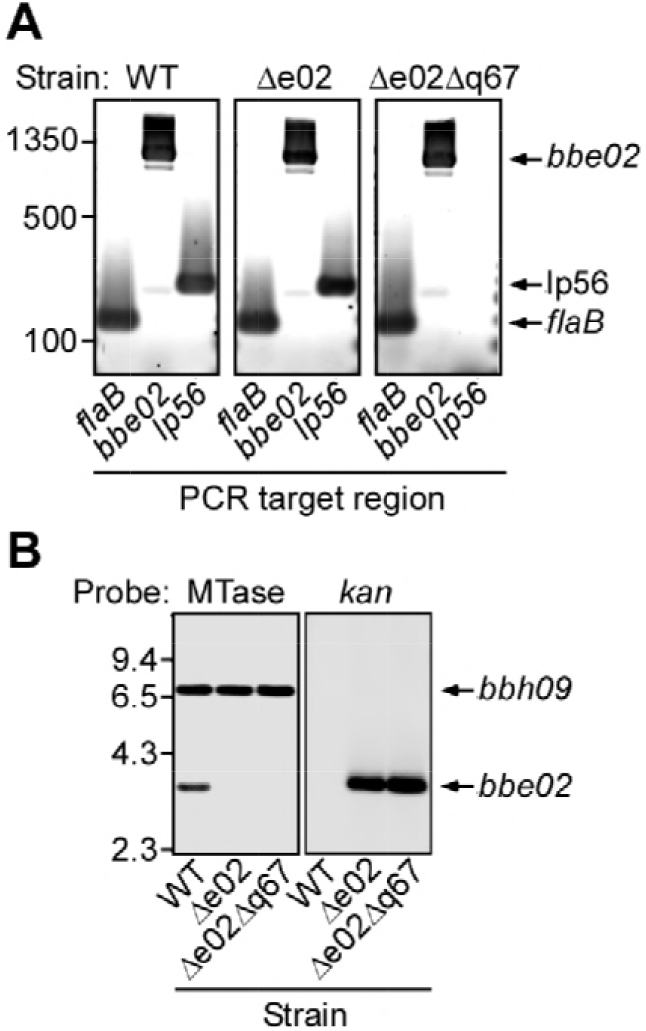
Confirmation of MTase gene disruption mutants. **A.** PCR analysis of total DNA from wild-type (WT) and deletion mutants as shown. Primer pairs for the chromosomal flaB gene were used as a positive control. All strains gave positive signal for the *bbe02* gene, likely due to nonspecific amplification of the homologous *bbh09* gene. As expected, no signal was detected for plasmid lp56 in the double MTase mutant. **B.** Southern blot analysis of NheI-digested plasmid DNA from wild-type and deletion mutant strains probed for *bbe02* (MTase) and *kan* as shown. Mutant strains were positive for the *kan* gene, and demonstrated the expected loss of signal for the *bbe02* gene, while the homologous *bbh09* gene was not affected. Location of size standards are shown to the left of the blots (bp for PCR, kbp for Southern blot), while expected sizes of the target fragments are shown on the right.

SMRT sequencing of genomic DNA from BbΔe02 and BbΔe02Δq67 revealed unambiguous changes in the profile of conserved m6A motifs for each strain (Table 2). BbΔe02 retained m6A modification at 95% of CGRKA motifs, however none of the other motifs found in wild-type *B. burgdorferi* were detected in this strain; identifying GNA<u>A</u>YG as the primary methylation target of BBE02, with DGDA<u>A</u>GG and DGGC<u>A</u>TG as promiscuous yet inefficient non-canonical target sites of this enzyme. No conserved m6A motifs were identified in BbΔe02Δq67, implicating CGRK<u>A</u> as the target site for methylation by BBQ67.

**Table 2.** Conserved m6A motifs in MTase mutants.

| Motif<sup>a</sup> | Type | # /<br>Genome | % Modified<sup>b</sup> |  |  |
| --- | --- | --- | --- | --- | --- |
|  |  |  | WT | Δe02 | Δe02Δq67 |
| CGRK<u>A</u> | m6A | 2,980 | 97 | 95 | NA |
| GNA<u>A</u>YG | m6A | 2,167 | 97 | NA | NA |
| DGDA<u>A</u>GG | m6A | 333 | 40 | NA | NA |
| DGGC<u>A</u>TG | m6A | 126 | 66 | NA | NA |

<sup>a</sup>Methylated adenine shown underlined<sup>b</sup>NA denotes motif not detected by SMRT analysis

Interestingly, no evidence was found of conserved m6A motifs in the absence of *bbe02* and *bbq67* in the given in vitro system, despite the presence of the intact *bbh09* gene and other loci with putative MTase motifs as predicted by ReBase (49), as well as evidence for an orphan MTase in relapsing fever Borrelia (50). A potential explanation for this phenomenon is that BBH09 is relevant exclusively during the enzootic cycle. It has been noted that although the predicted amino acid sequences of BBE02 and BBH09 share 92% similarity, the isoelectric points of these homologs are quite different (6.99 vs. 8.07), which may affect their activities (24). It is therefore feasible that standard in vitro culture conditions for *B. burgdorferi* favor the enzymatic activity of BBE02, but not BBH09. Likewise, sequence differences between BBH09 and BBE02 may reflect differences in substrate bases. It cannot be ruled out that BBH09 acts as a m5C methylatrasferase. SMRT sequencing is much less sensitive to detection of this type of DNA modification, and was therefore not examined in the current assay (36, 51). Finally, *bbh09* may not encode a functional MTase, and may be somehow required as an accessory molecule for BBE02 or BBQ67 functionality. RM enzymes typically function as dimers or tetramers, and often associate as heteromers (52). The requirement for accessory proteins for BBE02 and BBQ67 functionality is unlikely, however, given that clones lacking lp28-3 and other plasmids are not enriched during mutagenesis of *B. burgdorferi* as is often seen for lp25 and lp56 harboring the *bbe02* and *bbq67* genes (24, 25) (unpublished observations).

Taken together, the data presented here on methylation motif specificity and previous data on restriction protection demonstrate that *bbe02* and *bbq67* both encode bifunctional RM enzymes with structures similar to type IIC endonucleases, and MTase sequence specificities similar to the MmeI-family of type IIS enzymes (37). This family typically recognizes 6-7 nucleotide long contiguous sequence motifs, and modifies only a single strand for restriction protection. They possess both endonuclease and DNA methyltransferase activities in the same polypeptide and require AdoMet for endonuclease activity, making them members of the type IIG subgroup as well. Additionally, the restriction function of MmeI-family enzymes cleave ~20nt downstream of the recognition sequence, and require interaction between two molecules bound at specific recognition sites to achieve cutting. Although the BBE02 and BBQ67-specific motifs were not typically found in close proximity, it remains to be determined whether more distant DNA-protein complexes could associate to catalyze activity.

### Deletion of RM systems results in global changes in gene expression in *B. burgdorferi*

Although the primary function of RM systems in prokaryotes is thought to be restriction protection from foreign DNA, a recent study demonstrated global effects on gene expression by DNA methylation from RM systems (53). To examine this possibility in *B. burgdorferi*, transcriptome profiles were generated by deep sequencing of rRNA-depleted total RNA isolated from wild-type and the MTase mutants. In all, 1,317 annotated genes and ncRNAs (44) were detected. Using a false discovery rate (FDR) of ≤ 0.01, 417 genes were differentially expressed in BbΔe02 compared to wild-type, whereas 564 genes were differentially expressed in BbΔe02Δq67 (see tables S6 and S7 for complete expression profiles). Figures 5A-B show the number, direction, and fold-change of those genes differentially expressed in either BbΔe02 or BbΔe02Δq67, as well as the considerable overlap (305 genes) of those differentially expressed in both MTase mutants. Clusters of Orthologous Groups (COG) functional categories were all similarly represented in the differentially expressed genes lists, with the majority (58%) of differentially expressed genes as unclassified (data not shown). These data suggest that the DNA methylation patterns by both BBE02 and BBQ67 have distinct and wide-spread effects on gene expression in *B. burgdorferi*.

**Figure 5.**
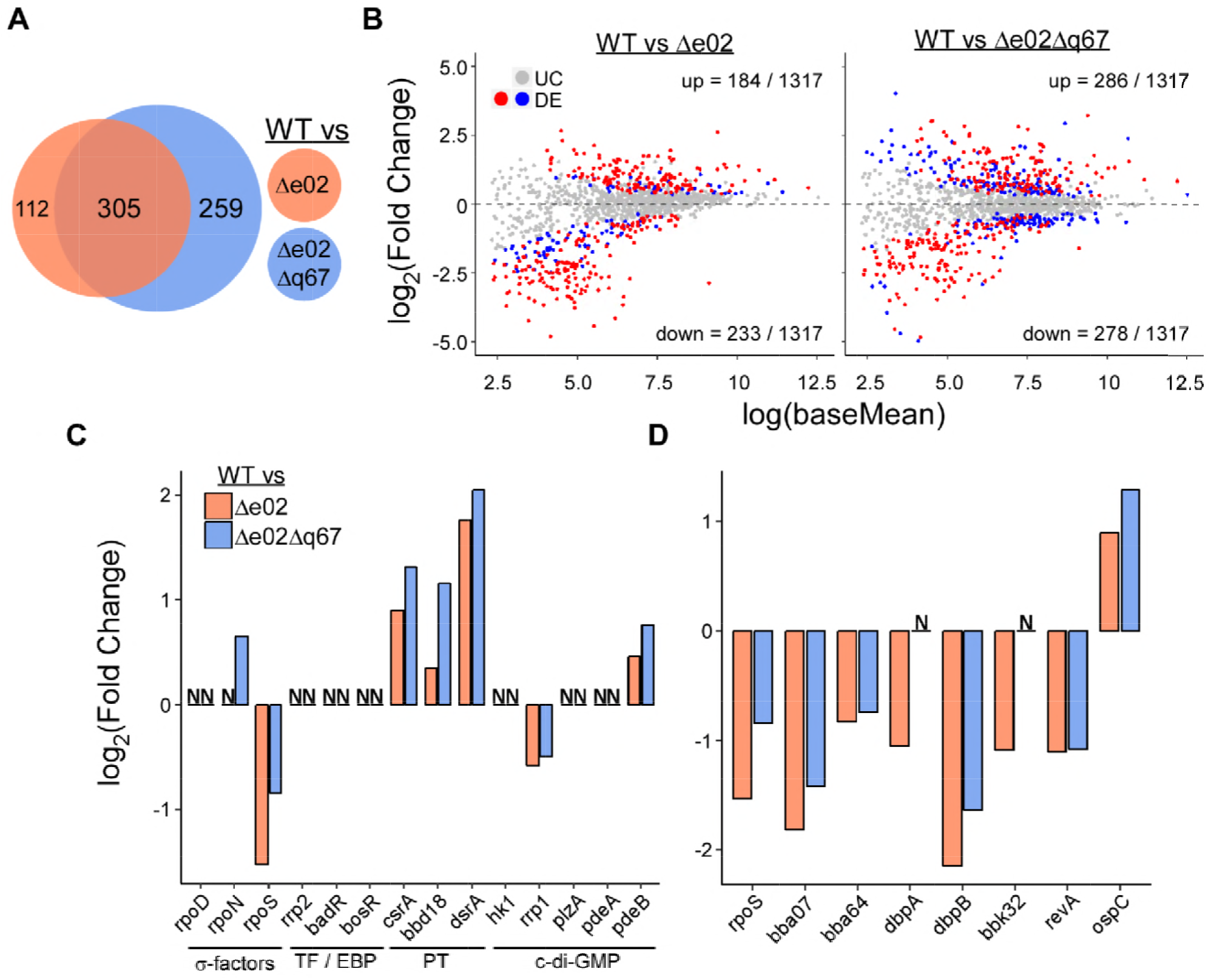
Deletion of MTases results in genome-wide changes in gene expression. **A.** Venn diagram showing the number of unique and shared differentially expressed genes (FDR ≤ 0.01) in MTase mutants compared to wild-type (WT) as determined by RNA-seq. **B.** Fold-change vs expression strength for all detectable genes. Red dots represent differentially expressed (“DE”) genes common to both MTase mutants, whereas blue dots represent genes differentially expressed in either BbΔe02 or BbΔe02Δq67 alone. Gray dots represent genes not significantly different between WT and MTase mutants (“UC”). Number of significant upregulated (“up”) and downregulated (“down”) genes are shown as proportions of all detectable genes. **C.** Differential expression of selected previously identified gene regulators. The alternative Sigma factor rpoS and post-transcriptional regulators of *rpoS* were most affected. “N” denotes no significant differential expression. Abbreviations: TF, transcription factor; EBP, enhancer-like binding protein; PT, post-transcriptional regulator; c-di-GMP, cyclic-di-GMP second messenger synthesis and/or effector pathway. **D.** Differential expression of *rpoS* and selected *rpoS*-dependent genes. All genes followed the expected pattern of differential expression based on *rpoS* expression with the exception of *ospC*, suggesting dysregulation by some additional regulatory network for that gene.

The enzootic cycle of *B. burgdorferi* requires transmission of the bacterium between animal hosts (predominantly small mammals) via ticks from the genus *Ixodes* (54). This dual-host lifestyle consisting of tick acquisition/colonization, transmission/acute mammalian infection, and persistent mammalian infection requires distinct gene expression patterns between these vastly different environments. Thus, much attention has been given to elucidating the regulators of adaptive gene expression profiles (55). Figure 5C shows the direction and magnitude of expression changes for selected genes involved in regulating gene expression in each MTase mutant. The housekeeping RNA polymerase sigma factor *rpoD* was unaffected in either mutant. In contrast, the mammalian-specific sigma factor *rpoS* was downregulated in both BbΔe02 and BbΔe02Δq67. This was coupled with a concomitant increase in expression of the known post-transcriptional *rpoS* repressors csrA, *bbd18*, and the small regulatory RNA *dsrA* (29, 56, 57). Likewise, several RpoS-dependent and “mammalian-specific” genes showed a concurrent downregulation including *bba07, bba64, dbpA, dbpB, bbk32*, and *revA* (58)(Figure 5D). Interestingly, the “canonical” RpoS-dependent gene *ospC* was upregulated in both MTase mutants. The *ospC* gene contains a unique operator region located immediately upstream of its RpoS-dependent promoter that is hypothesized to allow input from additional gene regulatory networks in order to independently repress *ospC* expression, while continuing to express the RpoS regulon during persistent infection (59). Thus, it is possible that any decrease in *ospC* expression in the MTase mutants due to downregulation of *rpoS* is offset by an even larger de-repression or activation by dysregulation of some yet unidentified regulatory network. Notably, the magnitude of *rpoS* downregulation was smaller in BbΔe02Δq67 than in BbΔe02. This pattern was consistent across all *rpoS*-dependent genes, including *ospC* (Figure 5D).

In addition to dysregulation of the *rpoS* regulon, both BbΔe02 and BbΔe02Δq67 showed a significant decrease in expression of the *rrp1* gene encoding a diguanylate cyclase response regulator (60) coupled with an increase in expression of the phosphodiesterase gene pdeB (61). Together, these data suggest the potential for decreased availability of cyclic-di-GMP within MTase mutants. Cyclic-di-GMP has been demonstrated to be important during tick acquisition and transmission of *B. burgdorferi*, and has been shown to have interplay with the *rpoS* regulon (62, 63). Collectively, these results demonstrate that gene regulatory networks relevant to survival during the enzootic cycle are influenced by endogenous RM systems in *B. burgdorferi*.

In order to examine the association between methylation events and disrupted gene expression profiles, the distribution of m6A motifs was compared between “unchanged” and “differentially expressed” genes for each mutant strain. No methylation profile was overrepresented in the list of differentially expressed genes compared to the proportion of all genes (Figure 6A). Additionaly, no association was observed when unchanged and differentially expressed genes were compared with respect to the presence/absence of m6A sites located immediately upstream of genes (Figure 6C), or with respect to the cumulative m6A distributions across/surrounding gene bodies (Figure 6D) or transcription start sites identified by Adams et al (64)(data not shown) using metagene analysis. Curiously, the median number of *bbe02* modification motifs was higher in those gene bodies where expression levels were unaffected compared to differentially expressed genes in both mutants (Figure 6B). It is likely that only a subset of m6A modifications have meaningful implications for gene expression changes, and those changes can be amplified through the affected gene regulatory networks. Therefore, global analysis of m6A distributions between unchanged and differentially expressed genes reveals little insight into the specific m6A changes responsible for altering the transcriptome. Similarly, Fang et al previously reported that although the expression of more than one-third of *E. coli* strain C227-11 genes were significantly altered when the PstI-like RM system, RM.EcoGIII, was deleted, there was not a compelling correlation detected between m6A modification events and differentially expressed genes (53).

**Figure 6.**
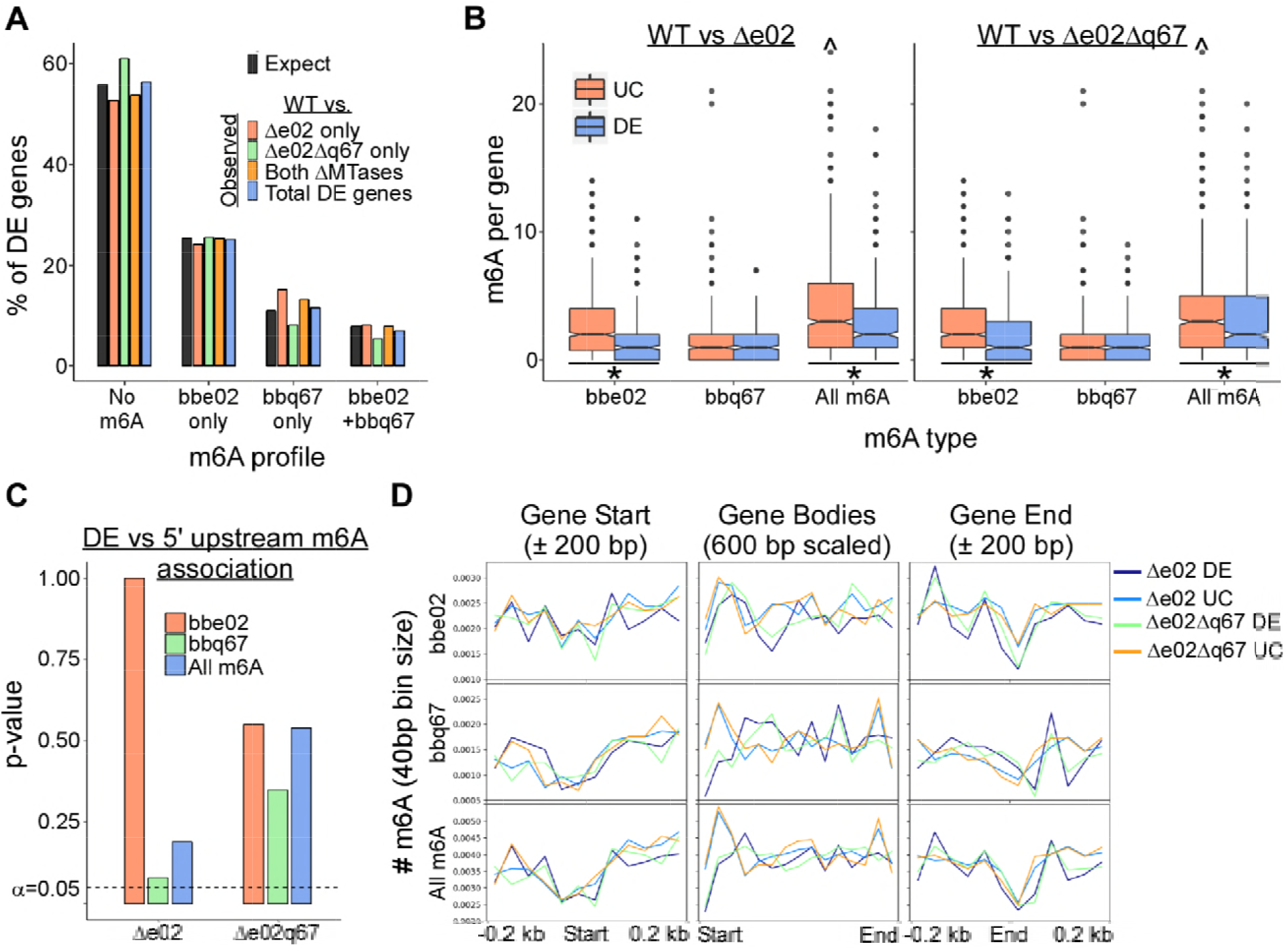
Correlation between m6A profile and differential expression. **A.** Proportion of genes differentially expressed (“DE”) in MTase mutants by methylation status. Black bars show the expected percentage based on the proportion of all gene bodies containing either no m6A motifs (“no m6A”), or at least one modification site for *bbe02* alone (“*bbe02* only”), *bbq67* alone (“*bbq67* only”), or at least one motif for both MTases (“*bbe02* + *bbq67*”). Colored bars show the observed percentage of differentially expressed genes with the corresponding m6A profiles as indicated in the legend. No methylation profile was overrepresented in the population of DE genes for either MTase mutant. **B.** Number of m6A sites per gene body by differential expression status for each MTase mutant. The number of *bbe02* motifs per gene was lower in differentially expressed (“DE”) genes compared to unchanged (“UC”) genes in both MTase mutants as determined by non overlapping 95% CIs and denoted by asterisks below the boxes. Carat character indicates outliers found beyond the Y-axis maximum. **C.** Association between the presence of at least one m6A modification located within 200 bp upstream of a gene start site and differential expression status. Fisher’s Exact test was performed on 2X2 contingency tables comparing gene counts for methylation status (+ or -) for each MTase with differential expression status (“DE” or “UC”). Barplot shows p-values from these tests for each m6A profile grouped by mutant strain. No association was below the α=0.05 threshold for significance, indicating a lack of association between upstream m6A and differential expression. **D.** Metagene analysis by differential expression status showing average m6A per 40 bp bin size across all gene bodies scaled to 600 bp (middle panel), as well as at gene start and end sites ± 200bp (left and right panels, respectively). Colored lines show comparisons of differentially expressed (“DE”) and unchanged (“UC”) genes in each mutant strain. All four groups showed similar profiles with no apparent change in the overall magnitude or significant shift from the gene ends.

One caveat of the differential gene expression data reported here is the potential for confounding effects from genetic differences other than MTase genes. Although the BbΔe02 strain is an isogenic deletion mutant of the wildtype strain, BbΔe02Δq67 lacks the entire lp56 plasmid harboring the *bbq67* gene as well as some other plasmid differences compared to the other two strains (Table S1). Although it cannot be ruled out that some of the gene expression differences observed in BbΔe02Δq67 are due to differences in the plasmid profile of this strain, it is likely that the vast majority of differentially expressed genes are a result of the lack of methylated DNA motifs for three reasons: 1) The differences in gene content in BbΔe02Δq67 do not include any predicted gene expression regulators, 2) the large numbers of affected genes in the isogenic BbΔe02 mutant demonstrate the global effects of differential genome methylation on gene expression, and 3) despite differences in clonal origin between the two muant strains, there is considerable overlap in the gene expression differences compared to the wildtype strain. Although this study clearly demonstrates the general phenomenon of altered gene expression resulting from differentially methylated genomes in *B. burgdorferi*, future investigations on specific individual differences would need to be verified experimentally.

Despite the global changes in gene expression observed in the MTase mutants when cultivated in vitro, it should be noted that *B. burgdorferi* lacking either *bbe02* alone or *bbe02* and *bbq67* are fully infectious in a laboratory murine model of infection as determined by ID50, joint swelling, and histopathology (26). These strains are also capable of completing the tick-mouse enzootic cycle under laboratory conditions, although lower pathogen burdens were reported in ticks for a strain lacking lp56 containing the *bbq67* gene (65). As the changes observed here for many genes including *rpoS* were relatively modest compared to the magnitude of induction in response to host-specific signals (66), it is possible that host signals are able to overcome gene dysregulation to levels sufficient for establishing infection in a laboratory setting. Nonetheless, loss of DNA methylation may have more subtle implications for gene regulation patterns and pathogen fitness during the enzootic cycle not detected in previous studies.

### Conclusions

The data presented here characterize for the first time the global m6A methylome during in vitro cultivation of *B. burgdorferi*, and assign specific methyltransferases responsible for all detectable conserved modification motifs. The MTases BBE02 and BBQ67 both methylate their primary target sequence at ~100% efficiency, consistent with restriction protection. Additionally, BBE02 appears to have inefficient, promiscuous, non-canonical activity at two additional sequence motifs that are not likely to provide restriction protection due to the much lower efficiency of methylation. It is not known whether these additional motifs carry biological significance, or if they are simply coincidental off-target methylation events. Nonetheless, as a consequence of the introduction of >5,000 methyl groups to the genome, methylation by both MTases has widespread effects on gene expression.

Homologues of the *bbe02* gene have been found in all identified *B. burgdorferi* isolates to date (25). In contrast, the presence of *bbq67* homologues and other predicted MTases varies between isolates. Rego et al hypothesized that these differences may act to drive strain heterogeneity in *B. burgdorferi*, and suggest a mechanism primarily related to restriction protection from horizontal gene transfer (25). The data presented here support alterations in gene expression profiles as another potential mechanism driving strain heterogeneity through RM diversification.

The data presented in this study have implications for laboratory models of Lyme disease research. The MTase genes *bbe02* and *bbq67* were first identified due to the high transformability of strains lacking these loci (24). As a consequence, many laboratories utilize MTase-deficient strains including the BbΔe02Δq67 strain used in this study (5A18-NP1) as surrogates for wild-type *B. burgdorferi* due to their ease of genetic manipulation (27-33). The findings reported here suggest that these strains may not be appropriate model organisms for *B. burgdorferi*, particularly for studies on gene regulation. There is a clear need for improvements to these model systems, including *B. burgdorferi*-specific cloning vectors devoid of recognition sites for cleavage by BBE02 or BBQ67, or mutant strains lacking the restriction endonuclease function of these RM enzymes while retaining MTase functions. Such tools would serve to overcome the low transformation efficiency of wild-type *B. burgdorferi*, while retaining the native methylome and gene expression profiles.

Overall, this study provides the first evidence of the functional consequences of RM systems in *B. burgdorferi* beyond restriction protection. Further studies involving strains with different profiles of RM systems, as well as biochemical characterizations of RM enzymes will further our understanding of the biology of this important group of human pathogens.

## EXPERIMENTAL PROCEDURES

### Bacterial strains and culture conditions

The *B. burgdorferi* strains A3 (wild-type) and 5A18-NP1 (BbΔe02Δq67), were kindly provided via gifts from Patti Rosa and Steven Norris, respectively. Both are clones of the sequenced B31 isolate (47), and their respective plasmid profiles have been previously described (26, 67)(Table S1). A3 was used to generate the mutant strain A3-NP1 (BbΔe02) as described. All *B. burgdorferi* clones were grown at 35°C under 5% CO2 in modified Barbour-Stoenner-Kelly medium (BSK) supplemented with 6% rabbit serum (68). Mutant strains were grown with kanamycin (200 μg/ml). Cell densities and growth phases were monitored by visualization under dark-field microscopy and counting using a Petroff-Hausser counting chamber.

### Generation of A3-NP1 mutant strain

In order to introduce the NP1 mutation into a wild-type A3 background, plasmid DNA was isolated from *B. burgdorferi* 5A18-NP1 using a Plasmid Midi Kit (Qiagen, Valencia, CA) for use as PCR template. A region encompassing the deleted portion of the *bbe02* gene containing an integrated kanamycin resistance cassette was amplified using primers P16 and P17 (26) (Table S8), and subsequently cloned into the vector pJET1.2 (Thermo Fisher Scientific, Waltham, MA). The resulting deletion plasmid was transformed into an *E. coli* intermediate strain for maintenance and propagation, and the insert was confirmed by restriction digest and DNA sequencing analysis.

A3 electrocompetent cells were cultivated and prepared as previously described (69). Cells were transformed with 50μg of the purified deletion construct described above, after which transformations were recovered for 24 hours in BSK followed by plating by limiting dilution with kanamycin selection to isolate clonal transformants. Deletion mutants were initially identified by PCR screening for the kanamycin resistance gene using primers P8 and P9 (70) (Table S8). Isolated plasmid DNA from kanamycin-positive clones was screened by PCR to determine endogenous plasmid content using primers specific for regions unique to each plasmid, as described previously (71). DNA from one representative clone containing all parental endogenous plasmids and the inserted kanamycin gene was selected for further analysis by Southern blot as described below.

### Southern blot hybridization

Plasmid DNA isolated from strains wild-type, BbΔe02, and BbΔe02Δq67 was digested with NheI (New England Biolabs, Ipswich, MA), and cleaned using PCR cleanup kit (Qiagen). Digested DNA was separated using 1% agarose gel electrophoresis followed by bidirectional transfer to two Amersham Hybond^™^ -N+ membranes for Southern blot analysis (GE Healthcare, Piscataway, NJ). Probes for *kan* and *bbe02* were generated by PCR from purified deletion construct plasmid or A3 plasmid DNA respectively, using primers P8 and P9 (*kan*) or P143 and P144 (*bbe02*) (Table S8) with the DIG Probe Synthesis Kit (Roche, Indianapolis, IN), according to the manufacturer’s instructions. Bands were detected using anti-DIG Fab fragment conjugated to alkaline phosphatase, and visualized using the chemiluminescent substrate CDP-Star (Roche).

### Genome sequencing and methylation analysis

DNA was isolated from a pool of three independent late log-phase cultures (5 mls per culture) for each *B. burgdorferi* strain by standard phenol-chloroform extraction, concentrated by precipitation with isopropanol, and cleaned using Agencourt AMPure XP beads (Beckman Coulter). Purified DNA was outsourced to the Deep Sequencing Core at the University of Massachusetts Medical School for library preparation and sequencing. Libraries were generated using a 20kb shear protocol, loaded onto one SMRT cell per sample, and sequenced on a PacBio RSII instrument using P6-C4 sequencing chemistry and a 360-minute data collection protocol.

The detection of modified bases and clustering of methylated sites to identify methylation-associated motifs was performed using the RS_Modification_and_Motif_analysis tool from the SMRT analysis package 2.3.0 (http://www.pacb.com/devnet/). Filtered subreads were aligned to the published *B. burgdorferi* B31 genome (see Table S3 for RefSeq numbers), and IPD ratios (observed/expect) were calculated using PacBio’s in silico kinetic reference computational control model. The accuracy of modification detection using this model was increased by comparing the observed IPD ratios to the expected signatures of the bacterial modification types m6A and m4C. Sequence motif cluster analysis was done using PacBio Motif finder v1 using the default quality value (QV) cutoff of 30.

Strand specificity and genomic distribution of modified bases were determined using BedOps v2.4.32 (72). To determine localized differences in motif modification, the genome was split into 1000bp segments with 250bp overlapping sequence. BED files of genomic modifications were intersected with genomic features and/or segments to obtain counting statistics. Metagene analysis of m6A distributions within genes was done using deepTools2 (73).

### RNA isolation and sequencing

Three independent cultures of each of wild-type and MTase mutant strains were grown to mid-late log phase (3 × 107 cells / ml) for RNA isolation. RNA was isolated by first adding 20 mls RNAprotect (Qiagen) to 10 mls culture to stabilize transcripts during processing. Cells were collected by centrifugation, resuspended in 1 ml of pre-warmed (65°C) Trizol reagent (Invitrogen), and frozen at −80°C overnight. Cell suspensions were thawed at room temperature, and RNA was isolated using the Direct-zol RNA mini kit (Zymo Research) following the manufacturers instructions. RNA concentration was determined using using a Qubit 2.0 Fluorometer (Life Technologies), and RNA integrity was verified using microfluidic-based capillary electrophoresis with an Agilent 2100 Bioanalyzer (RIN ≥ 9.8 for all samples).

Directional cDNA libraries were prepared from 5 μg of purified RNA as input using the ScriptSeq complete bacteria kit and SciptSeq index PCR primers (Illumina) following the manufacturer’s instructions. Libraries were analyzed on the Bioanalyzer to ensure appropriate size distributions. The cDNA library from one sample (Δe02) did not pass library QC, and was omitted. The remaining 8 indexed cDNA libraries were pooled and sequenced on two runs using the Illumina MiSeq using 150 cycle V3 kits (75bp PE reads were collected). Reads from each sample in the pool were demultiplexed via Illumina CASAVA software v1.8, and fastq files from the two sequencing runs were combined for each sample prior to analysis.

For analysis of RNA-seq data, adapters were removed from sequencing reads using Trimmomatic (74). Reads were aligned and counted using a transcriptome reference compiled from the *B. burgdorferi* genome (see Table S3 for RefSeq numbers) as well as the ncRNAs described by Arnold et al (44) using Salmon 0.81 (75). Differential expression analysis was conducted on the raw read counts using DEseq2 (76). Plasmids lp5, cp9, cp32-6, lp38, and lp56 were removed prior to differential expression analysis as they are missing in at least one of the strains analyzed.

### Data visualization

Data generated from DNA and RNA sequencing analysis was visualized using R v.3.3.0 (https://www.R-project.org/) with the following packages: sequence logos, ggseqlogo (77); circos plot, Rcircos (78); barplots, boxplots, and MA plots, ggplot2 (79).

### Sequence availability

Sequences have been deposited in the NCBI GEO sequence read archive database under ascession ### (sequences have been uploaded, however accession number not assigned yet).

## ACKNOWLEDGEMENTS

We thank Ali Divan for critical reading of this manuscript, and Hannah Ness for technical assistance. This work was funded by NIH/NIGMS P20GM113123-01 and NIH/NIGMS P20 GM104360-01 to C. A. Brissette, and NIH R03-AI113648 to B. Stevenson.

